# Prevalence of genetically similar *Flavobacterium columnare* phages across aquaculture environments reveals a strong potential for pathogen control

**DOI:** 10.1101/2020.09.23.309583

**Authors:** A Runtuvuori-Salmela, HMT Kunttu, E Laanto, GMF Almeida, K Mäkelä, M Middelboe, L-R Sundberg

## Abstract

Intensive aquaculture conditions expose fish to bacterial infections, leading to significant financial losses, extensive antibiotic use and risk of antibiotic resistance in target bacteria. *Flavobacterium columnare* causes columnaris disease in aquaculture worldwide. To develop a bacteriophage-based control of columnaris disease, we isolated and characterized 126 *F. columnare* strains and 63 phages against *F. columnare* from Finland and Sweden. Bacterial isolates were virulent on rainbow trout (*Oncorhynchus mykiss*) and fell into four previously described genetic groups A, C, E and G, with genetic groups C and E being the most virulent. Phage host range studied against a collection of 228 bacterial isolates demonstrated modular infection patterns based on host genetic group. Phages infected contemporary and previously isolated bacterial hosts, but bacteria isolated most recently were generally resistant to previously isolated phages. Despite large differences in geographical origin, isolation year or host range of the phages, whole genome sequencing of 56 phages showed high level of genetic similarity to previously isolated *F. columnare* phages (*Ficleduovirus, Myoviridae*). Altogether, this phage collection demonstrates a potential to be used in phage therapy.

**Significance Statement:** Bacteriophages were discovered already over a century ago, and used widely in treatment of bacterial diseases before the era of antibiotics. Due to harmful effects of antibiotic leakage into environment, aquaculture is a potential target for phage therapy. However, the development of efficient phage therapy approach requires detailed characterization of bacterial pathogen virulence and phage host range. Here, we describe phage-bacterium interactions in the fish pathogen *Flavobacterium columnare*. We found that genetically similar phages are found from different fish farms, and their infectivity cluster according to genetic group of bacteria. In addition, phages were able to infect bacterial hosts from other farms, which is a preferable trait considering phage therapy approach. However, the most recently isolated phages had broader host range than the previously isolated phages, suggesting a response in the phage community to evolution of resistance in the bacteria. These results show that designing phage therapy for aquaculture (and other) systems needs consideration of both temporal and geographical aspects of the phage-bacterium interaction.

## Introduction

During the past 20 years, aquaculture has been the fastest growing food production sector (FAO 2014), providing a source of protein for human consumption. Intensive aquaculture production is based on monocultures of certain species, which are reared in high population densities, resulting in increased transmission of infections (Pulkkinen *et al.*, 2010; Oidtmann *et al.*, 2011) and antibiotic use (FAO, 2014). Approximately 70 – 80 % of antibiotics in aquaculture may end up in the environment (Cabello *et al.*, 2013; Watts *et al.*, 2017), where they select for antibiotic resistance also in the environmental bacteria(Tamminen *et al.*, 2011; Yang *et al.*, 2013). The World Health Organization (WHO) has declared antibiotic resistance as a major risk for global health and food security, and means to control diseases without antibiotics are therefore urgently needed.

Due to increased issues with antibiotic resistance and lack of efficient vaccines, application of lytic bacteriophages (phages) has been suggested as an alternative for controlling pathogenic bacteria (reviewed by e.g. Watts, 2017; Kortright *et al.*, 2019). Their host specificity makes phages strong candidates as tools for targeted eradication of pathogenic bacteria. Indeed, use of phage as therapeutics has a long history in medicine (Almeida and Sundberg, 2020), and recently interest towards other types of applications has increased. Phages can be used, for example, to extend the food shelf life (Moye *et al.*, 2018), and are already in use against *Listeria* in salad, salmon and meat packages (Sulakvelidze, 2013; Lone *et al.*, 2016). Interest towards using phages in aquaculture has been steadily increasing the past decade, including studies on phage-bacterium interactions for important fish pathogens such as Flavobacteria, Vibrio, and Aeromonas (Castillo *et al.*, 2012; Silva *et al.*, 2014; Tan *et al.*, 2014).

*Flavobacterium columnare* (Bacteroidetes) is a Gram-negative bacterium (Bernardet *et al.*, 1996) causing columnaris epidemics at freshwater fish farms worldwide (Declercq *et al.*, 2013). Conditions such as water temperature over 18°C and high fish density promote columnaris epidemics, which spread rapidly in the rearing units, and may lead to high mortality if not treated by antibiotics (Suomalainen *et al.*, 2005; Karvonen *et al.*, 2010; Pulkkinen *et al.*, 2010). Moreover, the intensive production of rainbow trout has been shown to select for highly virulent *F. columnare* strains in the aquaculture environment (Sundberg *et al.*, 2016), with different genotypes being connected to different host species at the global scale (LaFrentz *et al.*, 2018).

However, how the pathogen population structure in aquaculture systems influences the genetic and phenotypic (especially host range) patterns in phages is not understood. Phages infecting *F. columnare* described so far have been relatively host specific with a narrow host range (Laanto *et al.*, 2011), and phage addition during experimental columnaris infection has shown promising results on the survival of rainbow trout fry (Laanto *et al.*, 2015; Gabriel M.F. Almeida *et al.*, 2019). However, for development of successful phage therapy approach, it is essential to obtain an overview of the diversity and spatial distribution of both phage and bacterial populations, and establish a phage collection that covers this diversity. This requires isolation, and subsequent genetic and functional characterization of phage and host communities, and description of the phage-host interactions.

In this study, we isolated new *F. columnare* strains and their phages from 10 different aquaculture locations in Finland and Sweden during summer 2017. Highly virulent bacterial strains occurred at several farms. Phage infection patterns were studied on 228 different bacterial strains isolated during 20013-2017. Bacterial isolates were genetically characterized and their virulence determined on rainbow trout (*Oncorhynchus mykiss*). Morphology of the phages was confirmed with transmission electron microscopy (TEM), and whole genome sequencing was performed for 56 of the isolated phages. We show that geographically distant *F. columnare* phages have very similar genomic composition and cluster according to the genetic groups of the host bacteria. The phages were able to infect bacteria isolated from different fish farms, however, the impaired capacity of phages isolated earlier to infect bacteria in a later time point suggests that bacteria evolve resistance against phage in the aquaculture conditions.

## Materials and Methods

### Isolation of bacteria

Samples from >10 fish farms were collected between June and August 2017 during columnaris disease outbreaks (Figure 1; exact number and locations of the Swedish fish farms are confidential and not known by the authors, so they are jointly marked as Farm 10). Water samples (1 000 ml) were collected from earthen ponds, fiberglass or plastic tanks, and from the outlet water of the farms. *F. columnare* was isolated in addition to water samples, directly from infected fish, using standard culture methods on Shieh agar plates supplemented with tobramycin (Song *et al.*, 1988; Decostere *et al.*, 1997). The obtained isolates were pure cultured, and stored at −80 °C with 10 % glycerol and 10 % fetal calf serum.

**Figure 1.**
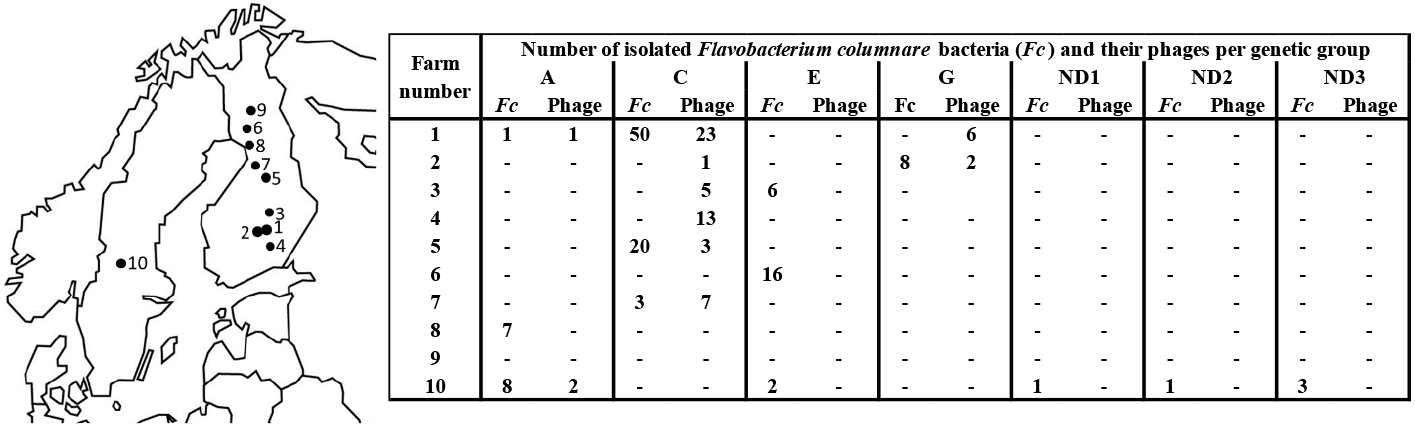
Sampling locations of fish farms in Finland and Sweden. On the left: Map of Northern Europe, where each number indicates a farm, where the water and fish samples were collected. Exact number and locations of Swedish farms are not known, and they are jointly marked as farm 10. On the right: Number of bacterial and bacteriophage isolates from individual fish farms. A, C, E, G, ND1, ND2 and ND3 indicate the different genetic groups of the isolated bacteria and the isolation hosts of the phage. ND = genetic group not determined.

### Genetic characterization of bacterial strains

Bacterial genomic DNA was extracted from overnight (o/n) bacterial liquid cultures with DNeasy® Blood & Tissue Kit (Qiagen) according to manufacturer’s instructions. For some bacterial isolates, a template for PCR reaction for genomovar and genetic group classification (see below) was obtained by picking one bacterial colony into 100 μl of sterile distilled water.

*F. columnare* isolates were classified to genomovars with restriction fragment length polymorphism (RFLP) analysis of 16S rRNA gene according to LaFrentz ((LaFrentz *et al.*, 2014) with some modifications. 16S rDNA was amplified by PCR (10 min at 95 °C; 40 cycles of 30 s at 94 °C, 45 s at 55 °C and 1 min at 72 °C; 10 min at 72 °C) using universal primers fD1 (Weisburg *et al.*, 1991) and 1500R (Triyanto and Hisatsugu Wakabayashi, 1999) with 1μM of each primer, 1 x DreamTaq Green buffer, 0,2 mM dNTP mix, 0,5 U Dream Taq DNA polymerase. PCR products (10 μl) were digested o/n at 37 °C with 3.3 U of *Hae*III. Restriction fragments were run in 12 % acrylamide gels, which were stained with ethidium bromide and visualized under UV-light.

The bacteria were further classified into genetic groups by RFLP of 16S – 23S ITS region correlating with the ARISA analysis designed for *F. columnare* (Suomalainen *et al.*, 2006). ITS region was amplified by PCR (2 min at 95 °C; 35 cycles of 30 s at 94 °C, 45 s at 52 °C and 3 min at 72 °C; 15 min at 72 °C) using primer pair rD1f (Weisburg *et al.*, 1991) and 23Sr (Borneman and Triplett, 1997) as above. PCR products (10 μl) were double-digested o/n at 37 °C with 3.3 U of both *Hae*III and *Hinf*I. Restriction fragments were run and visualized as described above.

### Virulence experiment

Thirty-four bacterial isolates representing all the genetic groups were selected for virulence testing on rainbow trout fry (Supplementary Table S1). Bacteria were revived from −80 °C by inoculation to 5 ml of Shieh medium and cultured overnight at 25 °C under constant agitation (120 rpm). Bacteria were enriched by subculturing (1:10) and incubating for 24 h. Bacterial cell density was measured as an optical density (OD, 595 nm; Multiscan FC Thermo Scientific) and colony forming units per ml (CFU/ml) estimated based on our previously determined OD-CFU relationship.

A total of 527 rainbow trout fry (*Oncorhynchus mykiss*), average weight 1.25 g, were randomly selected and placed individually into experimental aquaria with 500 ml of pre-aerated water (25 °C). For each bacterial isolate, 14-15 individual fish (20 for negative control) were infected by pipetting into each aquarium 500 μl of bacterial solution giving a final dose of 5×10^3^ CFU/ml. Shieh medium was used for negative controls. Fish morbidity and symptoms were observed one-hour intervals for 24 hours. Symptomatic fish not responding to stimuli were removed from the experiment and measured. To confirm the presence/absence of the bacterium, cultivations from gills of the dead fish were made on Shieh agar supplemented with tobramycin (Decostere *et al.*, 1997). At the end of the experiment, surviving fish were euthanatized with overdose of benzocaine. Mortality data was analyzed using Kaplan-Meier survival analysis in IBM-SPSS statistics 24 SPSS. High and medium virulence of individual isolates were defined by an estimated survival time of <15 h and > 15 hours, respectively, and low virulence was when no significant difference was detected compared to the control group.

Virulence test was performed according to the Finnish Act on Use of Animal for Experimental Purpose, under permission ESAVI/3940/04.10.07/2015 granted for Lotta-Riina Sundberg by the National Animal Experiment Board at the Regional State Administrative Agency for Southern Finland.

### Isolation and characterization of bacteriophages

For bacteriophage isolation, 500 ml of water sample was filtered using rapid flow filters (PES membrane, pore size 0.45 μm, Nalgene®). 5x Shieh medium was diluted to 1x using filtered water, and 1 ml of overnight-grown bacterial host (or mixture of hosts) was added (total volume 21 ml). In some isolations, Shieh was diluted to 0.5x Shieh supplemented with 0.1 % mucin (Gabriel M F Almeida *et al.*, 2019).

Four previously isolated and characterized *F. columnare* strains (genetic group in parenthesis) were used as enrichment hosts; B185 (G), B480 (E), B534 (A) and B537 (C). In addition, strains F514 (ND3), isolated from Sweden and FCO-F2 (C), isolated from Finland, were used in some of the enrichments (see Supplementary Table S1). All the strains were used both individually and as a mixture. In mixed cultures, the total bacterial cell density (at OD 595 nm) was adjusted to the same OD level as the bacterium with the lowest OD in the individual enrichments.

After incubating for 24 h at 25 °C (120 rpm), 0,5 ml samples were taken from enrichment cultures, centrifuged (3 minutes, 8000 x g), and supernatants collected. Presence of phages was detected using double layer agar method. 300 μl of fresh indicator bacterial culture and 300 μl of supernatant from the enrichment culture were mixed with 3 ml of soft Shieh agar (0.5 %) tempered to 46 °C, and poured on Shieh agar plates. When mucin was used in the isolation procedure, also the soft agar contained 0.1 % mucin. After one to two days of incubation at 25 °C, individual plaques were transferred to 500 μl Shieh medium, and subjected to three rounds of plaque purification.

Phage stocks were prepared by adding 5 ml Shieh medium on confluent and semi-confluent plates. Plates were incubated in a constant shaking (90 rpm) in cold room (7 °C) for approximately 12 h. The lysate was collected with a syringe, sterile-filtered (0.45 μm Acrodisc® Syringe Filters with Supor® Membrane), and stored at 7 °C for further use.

### Transmission electron microscopy

Three phages infecting different hosts and originating from different locations were selected for transmission electron microscopy (TEM) imaging. TEM samples were prepared from phage lysates on Cu-grids. A drop of lysate was added to the grid and after 15-30 seconds the grids were dried with moist filter paper (Whatman). Dried samples were incubated with phosphotungstic acid (1 % PTA, pH 7.5) for 30 – 60 seconds and dried as above. Grids were air dried o/n and protected from light. Imaging was done with JEOL JEM-1400 with 80 kVa, using LaB6 filament and program Radius.

### Phage host range

Host range of 73 phages (Table 1) was tested on 227 different bacterial hosts (Supplementary Tables S1, S2 and S3) using the double layer agar method. Two μl of each phage lysate and their 10- and 100-fold dilutions were spotted on bacterial lawns. Results were recorded after 2 days of incubation at room temperature. Phages were considered to infect the bacterium if all phage dilutions had clear spots or if individual plaques were observed. Turbid drop areas were considered as growth inhibition. Bacteria were considered resistant if the phage had no effect on the growth.

**Table 1.** The virulence (high, medium or low; based on estimated survival time, Kaplan-Meier survival Analysis) of 34 *Flavobacterium columnare* isolates on rainbow trout fry (*Oncorhynchus mykiss*) in a 24-hour experiment.

| Isolate | Isolation year | Country | Farm n:o | Genomovar | Genetic group | Mortality (%) | Mean estimated survival time (h) | Virulence |
| --- | --- | --- | --- | --- | --- | --- | --- | --- |
| FCO-F16 | 2017 | Finland | 3 | I | E | 100,0 | 12,87 | High |
| FCO-F81 | 2017 | Finland | 6 | I | E | 100,0 | 13,00 | High |
| FCO-F88 | 2017 | Finland | 6 | I | E | 100,0 | 13,07 | High |
| FCO-F2 | 2017 | Finland | 1 | I | C | 100,0 | 13,13 | High |
| FCO-F13 | 2017 | Finland | 3 | I | E | 100,0 | 13,53 | High |
| FCO-F118 | 2017 | Finland | 7 | I | C | 100,0 | 13,53 | High |
| FCO-F11 | 2017 | Finland | 3 | I | E | 100,0 | 13,60 | High |
| FCO-F22 | 2017 | Finland | 1 | I | C | 100,0 | 13,60 | High |
| FCO-F33 | 2017 | Finland | 1 | I | C | 100,0 | 13,67 | High |
| FCO-F58 | 2017 | Finland | 1 | I | C | 100,0 | 13,73 | High |
| FCO-F98 | 2017 | Finland | 5 | I | C | 100,0 | 13,80 | High |
| FCO-F50 | 2017 | Finland | 1 | I | C | 100,0 | 13,93 | High |
| FCO-F86 | 2017 | Finland | 6 | I | E | 100,0 | 14,00 | High |
| FCO-F30 | 2017 | Finland | 1 | I | C | 100,0 | 14,13 | High |
| FCO-F78 | 2017 | Finland | 5 | I | C | 100,0 | 14,20 | High |
| FCO-F41 | 2017 | Finland | 1 | I | C | 100,0 | 14,33 | High |
| F310 | 2000 | Sweden | 10 | I/II | ND3 | 100,0 | 14,93 | High |
| F387 | 2002 | Sweden | 10 | I | E | 93,3 | 15,33 | Medium |
| F383 | 2002 | Sweden | 10 | I/II | ND3 | 100,0 | 15,40 | Medium |
| FCO-F45 | 2017 | Finland | 8 | I | A | 100,0 | 15,53 | Medium |
| F514 | 2013 | Sweden | 10 | I/II | ND3 | 100,0 | 15,87 | Medium |
| FCO-F42 | 2017 | Finland | 8 | I | A | 100,0 | 15,87 | Medium |
| FCO-S1 | 2017 | Sweden | 10 | I | A | 100,0 | 16,00 | Medium |
| FCO-F32 | 2017 | Finland | 1 | I | A | 100,0 | 16,21 | Medium |
| FCO-F40 | 2017 | Finland | 1 | I | C | 100,0 | 16,27 | Medium |
| FCO-F47 | 2017 | Finland | 8 | I | A | 100,0 | 16,33 | Medium |
| 3/3449 | 2017 | Sweden | 10 | I | A | 100,0 | 16,40 | Medium |
| FCO-F5 | 2017 | Finland | 2 | I | G | 100,0 | 16,53 | Medium |
| FCO-F3 | 2017 | Finland | 2 | I | G | 100,0 | 16,60 | Medium |
| FCO-F9 | 2017 | Finland | 2 | I | G | 100,0 | 17,00 | Medium |
| FCO-F8 | 2017 | Finland | 2 | I | G | 100,0 | 17,13 | Medium |
| F524 | 2014 | Sweden | 10 | I | A | 100,0 | 19,87 | Medium |
| F512 | 2013 | Sweden | 10 | I | ND2 | 100,0 | 20,67 | Medium |
| F397 | 2002 | Sweden | 10 | I | ND1 | 20,0 | 22,33 | Low |
| Neg. control |  |  |  |  |  | 30,0 | 23,90 |  |

Interaction map of the phage-bacterium infection patterns was done with Gephi 0.9.2 (Bastian *et al.*, 2009) using the Force Atlas 2 network visualization algorithm (Jacomy *et al.*, 2014).

### Genetic characterization of bacteriophages

High-titer phage lysates were used for phage DNA extraction using the zinc chloride method modified from (Santos, 1991). DNase and RNase (final concentrations 1 μg/ml and 10 μg/ml, respectively) were added to filtered lysates and incubated at 37 °C for 30 minutes. Filtered ZnCl_2_ (final concentration 0.2 M) was used. Proteinase K final concentration was 0.8 mg/μl (Thermo Scientific). DNA was purified and extracted using phenol:chloroform:isoamyl alcohol (25:24:1) (Sambrook and Russell, 2001) and precipitated with isopropanol. DNA pellets were dissolved to 10-30 μl of dH_2_O. Purity of DNA (absence/presence of host DNA) was checked with PCR using universal bacterial 16S primers fD1 and rD1 (Weisburg *et al.*, 1991).

56 phage genomes were sequenced at the Institute for Molecular Medicine Finland (FIMM) by Illumina (HiSeq). Phage genomes were assembled by mapping the reads to reference sequence (KY979242 for genotype C-infecting phages, and NC_027125 for genotype G-infecting phages) using Geneious assembles (Geneious 7.1.4 and later versions). Genotype A-infecting phages were *de novo* assembled with Velvet 1.2.10 (in Geneious created by Biomatters Ltd). Genome ends were checked from V186 and S1 with Sanger sequencing using primers designed for *F. columnare* phage FCV-1 genome end verification (Laanto *et al*., 2017).

Open reading frames (ORFs) were predicted using GenemarkS (Besemer *et al.*, 2001) and Glimmer (Kelley *et al.*, 2012). Blastp (Jacob *et al.*, 2008) and HHPred (Söding *et al.*, 2005) were employed for annotating the putative function of the ORFs. Genomes were aligned using MUSCLE (Edgar, 2004)using default settings suggested by Geneious 7.1.4 (Biomatters Ltd.). Phage V182 isolate from farm 1 in 2014 was included in alignments as a reference to the last time point of the phage genome evolution dataset published earlier (Laanto et al., 2017). Genome comparison of the three phages infecting different host genotypes was created with Easyfig (Sullivan *et al.*, 2011) employing BLASTX.

Phage genetic distances were analysed with Victor (Meier-Kolthoff and Göker, 2017). All pairwise comparisons of the nucleotide sequences were conducted using the Genome-BLAST Distance Phylogeny (GBDP) method (Meier-Kolthoff *et al.*, 2013) under settings recommended for prokaryotic viruses (Meier-Kolthoff and Göker, 2017).

The resulting intergenomic distances were used to infer a balanced minimum evolution tree with branch support via FASTME including SPR postprocessing (Lefort *et al.*, 2015) for each of the formulas D0, D4 and D6, respectively. Branch support was inferred from 100 pseudo-bootstrap replicates each. Trees were rooted at the midpoint (Farris, 1972) and visualized with FigTree (Rambaut, 2016). Taxon boundaries at the species, genus and family level were estimated with the OPTSIL program (Göker *et al.*, 2009), the recommended clustering thresholds (Meier-Kolthoff and Göker, 2017) and an F value (fraction of links required for cluster fusion) of 0.5 (Meier-Kolthoff *et al.*, 2014).

## Results

### Isolation and genetic characterization of *F. columnare* strains

We isolated 111 *F. columnare* strains from water samples collected from 10 different locations in Finland and Sweden. In addition, 15 Swedish *F. columnare* strains were obtained from National Veterinary Institute, Sweden (Supplementary Table S1). Globally, *F. columnare* strains can be classified into six genomovars (I, I/II, II, II-A, II-B and III) by restriction fragment length polymorphism of 16S rDNA (Triyanto & Wakabayashi, 1999; LaFrentz et al., 2014, García et al., 2018). In this study, all isolated bacterial strains were classified as genomovar I strains, except for the previously isolated Swedish strains F310, F383 and F514, which were classified as genomovar I/II (Supplementary Table S1).

From altogether 126 bacterial isolates, 121 Finnish and Swedish strains could be assigned into previously defined genetic groups (A, C, G, E, (Suomalainen *et al.*, 2006) using RFLP analysis of the of internal transcribed spacer (ITS) region between 16S rRNA and 23S rRNA genes. Most of the strains fell into genetic group C (73 strains) or E (24 strains) (Figure 1; Supplementary Table S1), whereas eight strains belonged to group G and 16 to group A. Five Swedish strains could not be assigned to any of the previously defined genetic groups, and were designated as ND1, ND2 and ND3.

In general, the isolates from each fish farm represented a specific genomic group (Figure 1) suggesting that specific genomic groups are dominating the *F. columnare* communities at the individual fish farm. Only from the most frequently sampled farm (Farm 1), we isolated bacteria belonging to two genetic groups (A and C) (Figure 1; Supplementary Table 1). *F. columnare* was not isolated from farms 4 and 9.

### Virulence experiment

Virulence of 34 selected *F. columnare* isolates representing different genetic groups was studied on rainbow trout fry. Of the isolates, 17 were categorized as high virulence (estimated survival time <15 h), 16 as medium virulence (estimated survival time >15 h), and one as low virulence (no difference to uninfected control: *p* = 0.662, Kaplan-Meier survival analysis) isolate. The virulence observed among isolates belonging to genetic groups E and C (Table 1), were significantly higher than for group A and G isolates (p < 0.001) (Figure 2), with significanty faster cumulative mortality caused by group E than group C isolates (p< 0.001). Mortality caused by genetic groups A and G, on the other hand, did not differ from each other (*p* = 0.865).

**Figure 2.**
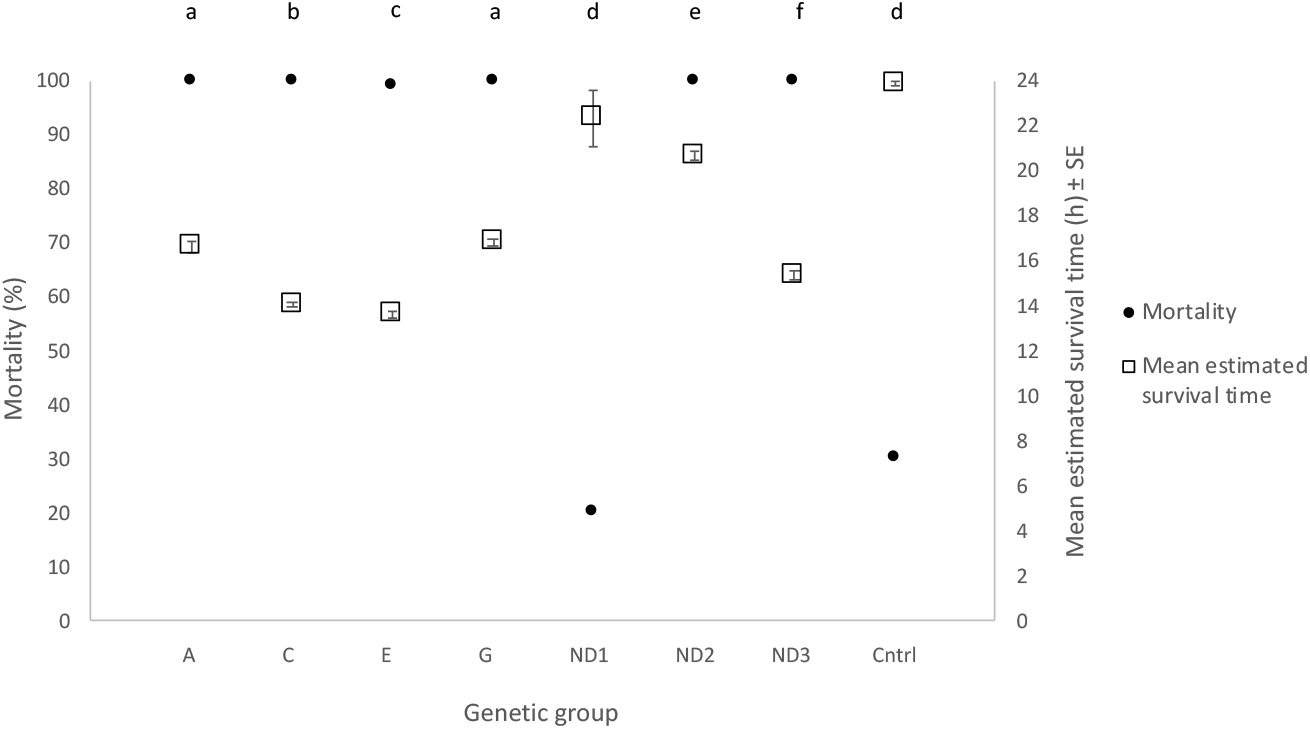
Mortality percent and mean estimated survival time (± SE) of rainbow trout (*Oncorhynchus mykiss*) during 24-h experimental infection with *Flavobacterium columnare* isolates representing genetic groups A, C, E and G. ND = genetic group not determined, Cntrl = control with no bacterial infection. Different lower-case letters indicate statistical difference in cumulative mortality (Kaplan-Meier Survival Analysis) between the genetic groups.

In addition to the genetic group, initial isolation source of the bacterium (rearing tank water or fish, Supplementary Table S1) had a significant effect on bacterial virulence, with isolates from tank water being more virulent than bacteria isolated from fish (*p* < 0.001). There were also differences in mortalities caused by bacteria isolated from different fish species/rearing tanks of different species (*p* < 0.001), the isolates from salmon being the most and the isolates from trout the least virulent.

### Characterization of bacteriophages

Sixty-three bacteriophages were isolated from the water samples originating from seven different fish farms (Figure 1; Table 2). The majority (52 out of 63) of the phages were isolated against hosts from genetic group C, while the rest of the phages were isolated against genetic groups A or G. No phages were isolated from mixed bacterial enrichment cultures or with genetic group E bacteria. Phages infecting hosts from different genetic groups were morphologically similar under TEM and displayed typical characteristics of myoviruses, icosahedral capsid with a non-bending tail (Supplementary Figure S1). Interestingly, phages against genetic group C were isolated from fish farm 4, even though no *F. columnare* was detected from the farm during the sampling.

**Table 2.** Bacteriophages isolated in this study. Phages marked with asterisks have been isolated and characterized in previous studies. ORFs were predicted with GenemarkS. For details of isolation hosts, see Supplementary Table S2.

| Phage | Fish farm n:o | Isolation year/date | Source | Isolation host /genetic group | Number of predicted ORFs | Genome length kbp | Accession number | Genetically identical isolates |
| --- | --- | --- | --- | --- | --- | --- | --- | --- |
| FCL-2* | 2 | 2008 | - | B185/G |  | 47,1 | NC_027125 |  |
| FCV-1* | 1 | 2009 | - | C1/C |  | 46,5 | NC_041845 |  |
| V182* | 1 | 2014 | - | B245/C |  | 49,1 | KY979242 |  |
| V183 | 1 | 2015 | - | B245/C | 76 | 49,1 | MT585311 | g |
| V184 | 1 | 2015 | - | B245/C | 76 | 49,1 | MT585312 |  |
| V186 | 1 | 2015 | - | B067/A | 74 | 46,5 | MT585313 |  |
| V188 | 1 | 2015 | - | B245/C | 76 | 49,1 | MT585314 |  |
| V189 | 1 | 2015 | - | B245/C | 76 | 49,1 | MT585315 |  |
| FCOV-S1 | 10 | 3.8.2017 | Tank water | B534/A | 74 | 46,5 | MK756094 | a |
| FCOV-S2 | 10 | 3.8.2017 | Tank water | B067/A | 74 | 46,4 | MK756095 | a |
| FCOV-F1 | 1 | 6.7.2017 | Tank water | B537/C | 76 | 49,1 | MT585273 | b |
| FCOV-F2 | 1 | 6.7.2017 | Tank water | B537/C | 76 | 49,1 | MK756083 | b |
| FCOV-F3 | 1 | 6.7.2017 | Tank water | B537/C | 76 | 49,1 | MT585274 | b |
| FCOV-F4 | 1 | 6.7.2017 | Tank water | B537/C | 76 | 49,1 | MT585275 | b |
| FCOV-F5 | 3 | 24.7.2017 | Tank water | B537/C |  |  |  |  |
| FCOV-F6 | 3 | 24.7.2017 | Tank water | B537/C | 76 | 49,1 | MK756084 | b |
| FCOV-F7 | 3 | 24.7.2017 | Tank water | B537/C | 76 | 49,1 | MT585276 | b |
| FCOV-F8 | 1 | 7.8.2017 | Farm outlet water | B537/C | 76 | 49,1 | MT585277 | c |
| FCOV-F9 | 1 | 7.8.2017 | Tank water | B537/C | 76 | 49,1 | MK756085 | c |
| FCOV-F10 | 1 | 7.8.2017 | Tank water | FCO-F2/C | 76 | 49,1 | MT585278 | b |
| FCOV-F11 | 1 | 7.8.2017 | Tank water | FCO-F2 | 76 | 49,1 | MT585279 | b |
| FCOV-F12 | 4 | 8.8.2017 | Farm outlet water | FCO-F2/C | 76 | 49,1 | MT585280 | e |
| FCOV-F13 | 1 | 7.8.2017 | Tank water | B185/G | 74 | 47,2 | MK756086 |  |
| FCOV-F14 | 1 | 7.8.2017 | Tank water | B185/G | 74 | 47,2 | MT585281 |  |
| FCOV-F15 | 1 | 7.8.2017 | Tank water | B185/G | 74 | 47,2 | MT585282 |  |
| FCOV-F16 | 1 | 7.8.2017 | Tank water | B185/G | 74 | 47,2 | MK756087 |  |
| FCOV-F17 | 1 | 7.8.2017 | Tank water | B537/C | 76 | 49,1 | MT585283 | c |
| FCOV-F18 | 1 | 7.8.2017 | Tank water | FCO-F2/C | 76 | 49,1 | MT585284 | c |
| FCOV-F19 | 7 | 23.8.2017 | Tank water | B537/C | 76 | 49,1 | MT585285 | f |
| FCOV-F20 | 5 | 18.8.2017 | Tank water | B537/C | 76 | 49,1 | MT585286 | f |
| FCOV-F21 | 5 | 18.8.2017 | Tank water | B537/C | 76 | 49,1 | MT585287 | f |
| FCOV-F22 | 7 | 23.8.2017 | Farm outlet water | FCO-F2/C | 76 | 49,1 | MT585288 | f |
| FCOV-F23 | 1 | 7.8.2017 | Tank water | B537/C | 76 | 49,1 | MT585289 | c |
| FCOV-F24 | 1 | 7.8.2017 | Tank water | FCO-F2/C | 76 | 49,1 | MT585290 | b |
| FCOV-F25 | 1 | 7.8.2017 | Tank water | B537/C | 76 | 49,1 | MK756088 | b |
| FCOV-F26 | 1 | 7.8.2017 | Tank water | FCO-F2/C | 76 | 49,1 | MT585291 | b |
| FCOV-F27 | 1 | 7.8.2017 | Tank water | FCO-F2/C | 76 | 49,2 | MK756089 | b |
| FCOV-F28 | 1 | 7.8.2017 | Tank water | B537/C | 76 | 49,1 | MT585292 |  |
| FCOV-F29 | 1 | 7.8.2017 | Tank water | B537/C | 76 | 49,1 | MT585293 | d |
| FCOV-F30 | 1 | 7.8.2017 | Tank water | B537/C | 76 | 49,1 | MT585294 | c |
| FCOV-F31 | 1 | 7.8.2017 | Tank water | FCO-F2/C | 76 | 49,1 | MT585295 | d |
| FCOV-F32 | 1 | 7.8.2017 | Tank water | FCO-F2/C | 76 | 49,1 | MT585296 | b |
| FCOV-F33 | 4 | 8.8.2017 | Tank water | B537/C | 76 | 49,1 | MT585297 | e |
| FCOV-F34 | 4 | 8.8.2017 | Tank water | B537/C | 76 | 49,1 | MT585298 | e |
| FCOV-F35 | 4 | 8.8.2017 | Tank water | B537/C | 76 | 49,1 | MT585299 | e |
| FCOV-F36 | 4 | 8.8.2017 | Tank water | B537/C | 76 | 49,1 | MT585300 | e |
| FCOV-F37 | 4 | 8.8.2017 | Tank water | B537/C | 76 | 49,1 | MK756090 | e |
| FCOV-F38 | 4 | 8.8.2017 | Tank water | B537/C | 76 | 49,1 | MT585301 | e |
| FCOV-F39 | 4 | 8.8.2017 | Tank water | FCO-F2/C | 76 | 49,1 | MT585302 | e |
| FCOV-F40 | 4 | 8.8.2017 | Tank water | FCO-F2/C | 76 | 49,1 | MT585303 | e |
| FCOV-F41 | 4 | 8.8.2017 | Tank water | FCO-F2/C | 76 | 49,1 | MT585304 | e |
| FCOV-F42 | 4 | 8.8.2017 | Tank water | FCO-F2/C | 76 | 49,1 | MT585305 | e |
| FCOV-F43 | 4 | 8.8.2017 | Tank water | FCO-F2/C | 76 | 49,1 | MT585306 | e |
| FCOV-F44 | 4 | 8.8.2017 | Tank water | FCO-F2/C | 76 | 49,1 | MT585307 | e |
| FCOV-F45 | 2 | 15.8.2017 | Tank water | B185/G | 74 | 47,2 | MK756091 |  |
| FCOV-F46§ | 2 | 28.8.2017 | Tank water | B185/G | 74 | 47,2 | MK756092 |  |
| FCOV-F47§ | 5 | 18.8.2017 | Tank water | B537/C | 76 | 49,1 | MT585308 | f |
| FCOV-F48§ | 2 | 15.8.2017 | Tank water | B537/C | 76 | 49,1 | MT585309 | g |
| FCOV-F49§ | 7 | 23.8.2017 | Tank water | B537/C |  |  |  |  |
| FCOV-F50§ | 7 | 23.8.2017 | Tank water | B537/C |  |  |  |  |
| FCOV-F51§ | 3 | 24.7.2017 | Tank water | B537/C |  |  |  |  |
| FCOV-F52§ | 7 | 7.8.2017 | Tank water | B537/C |  |  |  |  |
| FCOV-F53§ | 7 | 7.8.2017 | Tank water | B537/C |  |  |  |  |
| FCOV-F54§ | 1 | 3.9.2017 | Tank water | B185/G | 74 | 47,2 | MK756093 |  |
| FCOV-F55§ | 1 | 3.9.2017 | Tank water | B185/G |  |  |  |  |
| FCOV-F56§ | 1 | 3.9.2017 | Tank water | B067A | 74 | 46,5 | MT585310 |  |
| FCOV-F58§ | 7 | 7.8.2017 | Tank water | B537/C |  |  |  |  |
| FCOV-F59§ | 1 | 7.8.2017 | Tank water | B537/C |  |  |  |  |
| FCOV-F60§ | 1 | 7.8.2017 | Tank water | B537/C |  |  |  |  |
| FCOV-F61§ | 1 | 7.8.2017 | Tank water | B537/C |  |  |  |  |
| FCOV-F62§ | 3 | 24.7.2017 | Tank water | B537/C |  |  |  |  |

\* phages characterized in (Laanto *et al.*, 2011; Laanto *et al.*, 2017)
§ phages isolated with mucin supplement (Gabriel M.F. Almeida *et al.*, 2019)

Host range of *F. columnare* phages isolated in this study and previously isolated phages (Table 2) was tested in total with 227 different bacterial strains (Supplementary Table S4). 36 Finnish and 8 Swedish new bacterial strains were susceptible to one or more of the phages. In cases where clear infections and plaques were not observed, majority of the phages caused growth inhibition on bacterial lawn (Supplementary Table S4).

The phages isolated in 2017 infected both contemporary and previously isolated *F. columnare* strains within the genetic cluster, regardless of the isolation origin of the bacteria. Network analysis of the phage infection patterns revealed clustered interactions defined by the genetic host genetic group (Figure 3, see also Supplementary Table S4). In other words, phages isolated against a host from a specific genetic group, infected only bacteria from the same genetic group, with few exceptions connecting the clusters (below). In cluster C, the pattern differentiated also by isolation year, as the phages isolated earlier (V183 and V184, V188, V189 isolated in 2015 from farm 1) were able to infect only bacterial strains isolated earlier (B245 in 2009 and B526 in 2012).

**Figure 3.**
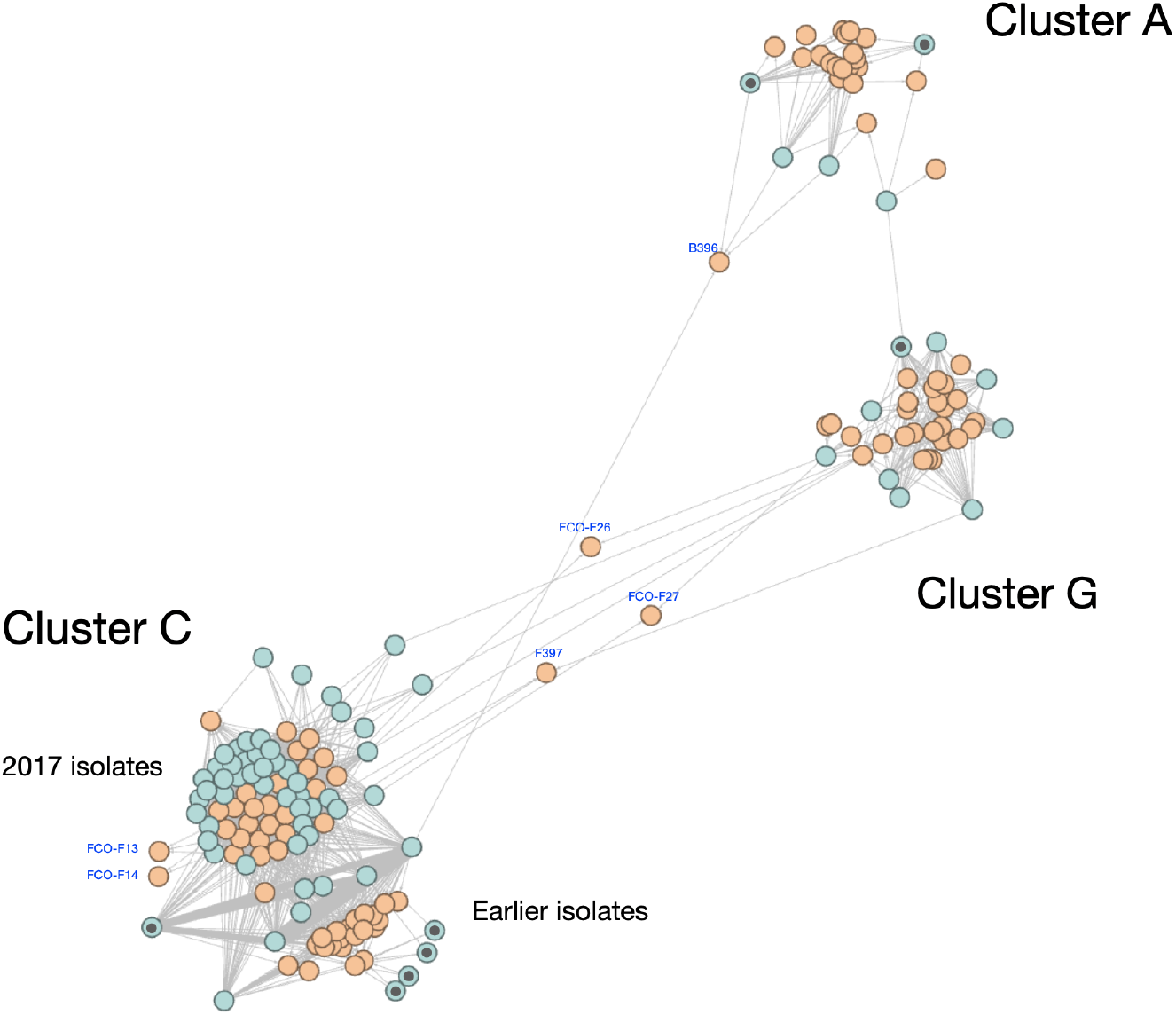
Phage-bacterium interaction network. Infection patterns of phages (light blue) in bacterial hosts (orange) are clustered based on the genetic group of the bacterial host (A, C, and G). Dark dot within phage circles indicate phages isolated earlier. Some key bacterial species infected by two different phage groups (FCO-F26, FCO-F27, F397, B396) or belonging to genetic group E (FCO-F13 and 14) are indicated. Network was visualized using Force Atlas 2 algorithm in Gephi, where modularity of the community is interpreted by comparing the nodes with each others.

Bacteria isolated in 2017 were generally resistant to phages isolated earlier (Figure 3, Supplementary Table S4). The phages characterized in this study did not infect bacteria isolated in USA, or other tested Flavobacterium species.

Few phages deviated from these cluster patterns, showing cross-infection to hosts from a different genetic group. Four phages isolated with G host B185 (FCOV-F13, F14, F15 and F16) were able to also infect genetic group A (3/3449, 5/3451, isolated from Sweden in 2017) bacteria (Figure 5, Supplementary Table S4). In addition, FCOV-F16 was able to infect also ND1 (isolate F397) and ND3 (isolate F310) strains from Sweden. C-genotype infecting phages FCOV-F25, F26, and F27 were able to infect bacteria isolates from genetic groups E (isolates FCO-F13 and FCO-F14), and G (isolate B442). FCOV-F28 was able to infect bacteria from genetic group G (isolate B442) in addition to group C bacteria (Supplementary Table S4). Interestingly, these extended host ranges were associated with minimal or no genetic differences in phage genomes (see below)

**Figure 5.**
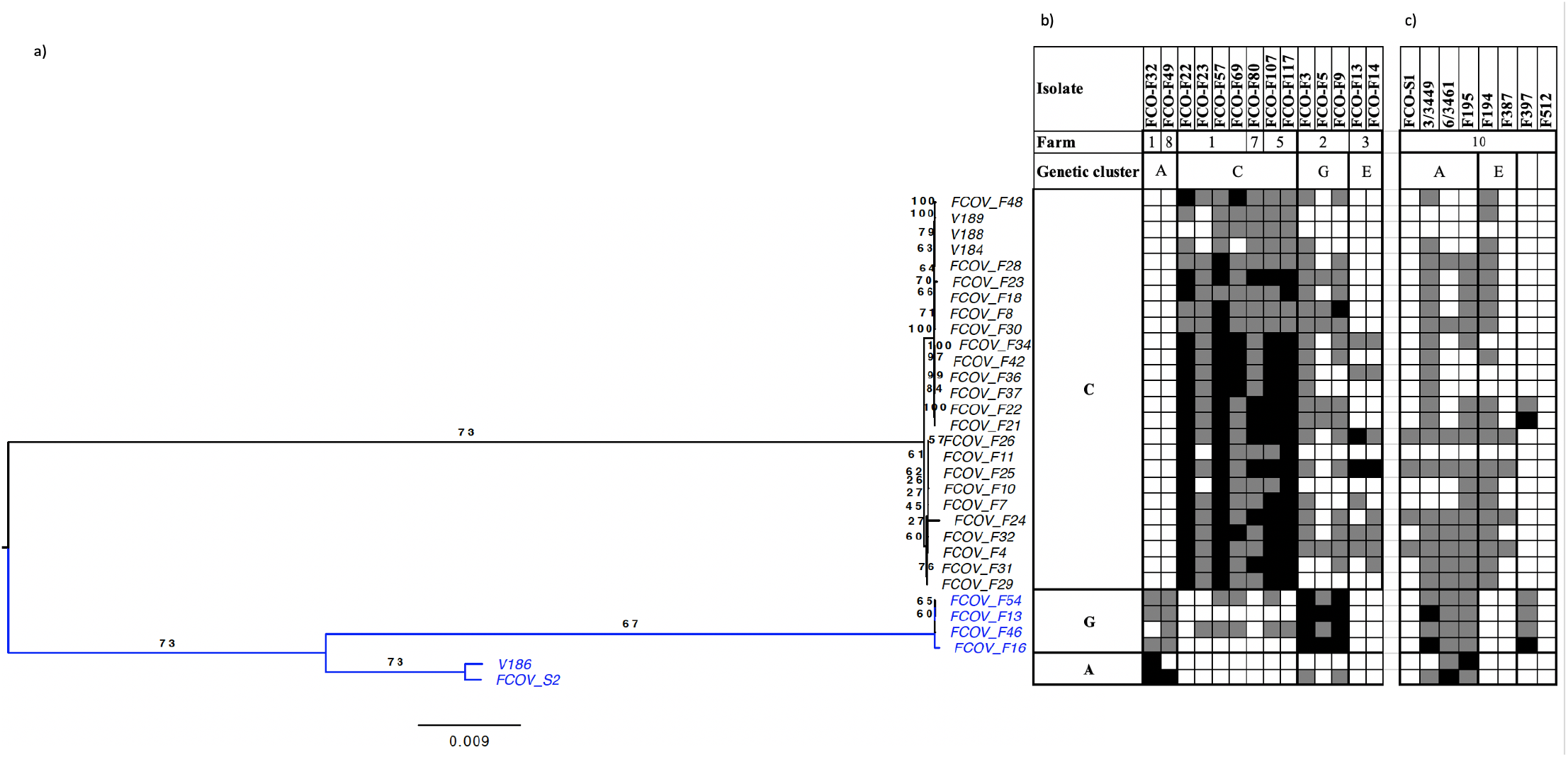
Genetic distance versus host range of sequenced phages. **A)** Genome BLAST Distance Phylogeny (GBDP) tree. The numbers above branches are GBDP pseudo-bootstrap support values from 100 replications. Host range of sequenced bacteriophages against representative bacterial strains isolated in **B)** Finland, and **C)** in Sweden. Black square indicates phage infection, grey square growth inhibition and white square bacterial resistance. Complete host range is provided in Supplementary materials.

### Genetic characterization of bacteriophages

Sequencing of the phage genomes revealed highly similar genomes despite the different host range of the phages (Figure 4). Comparing phage genomes across infection clusters (i.e. across host genomic groups), showed a nucleotide level identity between phage genomes of C- and G-phages of 82 % whereas A and G phages shared 91 % identity, and A and C phages 87% Figure 4).

**Figure 4.**
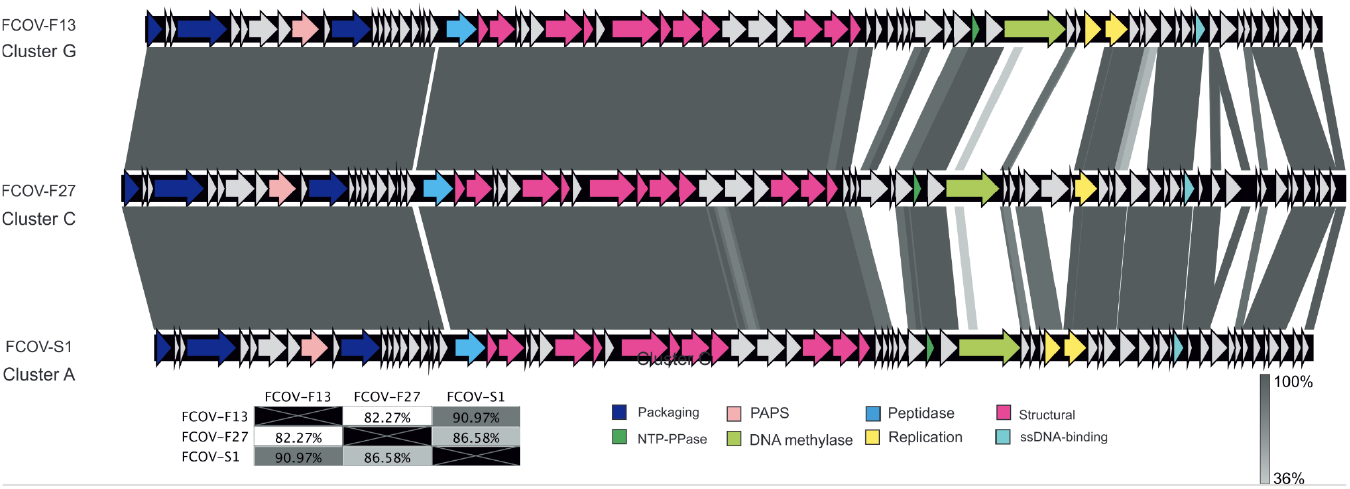
Genomic comparison of representative phages of the Clusters A (FCOV-S1), C (FCOV-F27) and G (FCOV-F13). Arrows in image indicate locations and orientations of ORFs. ORFs with unknown function are marked with grey, ORFs with putative function are marked with colors indicating the putative function as marked in the bottom. PAPS stands for 3’-phosphoadenosine-5’-phosphosulfate. The nucleotide level identity between the genomes in indicated in the box bottom left. Gray regions between the genomes indicate the level of identity (see the legend).

Length of the linear phage genomes varied from 46 kbp with 74 ORFs (Cluster A phages) and 47 kbp with 74 ORFs (Cluster G) up to 49 kbp with 76 ORFs (Cluster C) (Supplementary Table S2). Most of the differences between phages infecting A-, C- and G-hosts were located at the end of the genome, whereas approximately the first 29 kbps of the genomes were more conserved (Figure 4, Supplementary Table S5). The conserved area consists of the predicted e.g. the packaging and structural genes. Genes related to lysogeny, bacterial virulence or antibiotic resistance were not found.

Phylogenetic tree based on complete phage genomes indicated similar clustering as the phage infection patters (Figure 5). Phylogenomic GBDP tree inferred using the formula D6 yielded average support of 70%. The OPTSIL clustering yielded two species clusters (C-phages and G+A phages), and one genus level. All the phages characterized here can be assigned to the unclassified Ficleduoviruses (*Myoviridae*).

Within the individual infection and species clusters, the phages had very similar genomes. The nucleotide level identities between phage genomes within infection clusters A, C and G (Figure 3) were high: 99.7 % – 100 % between A-phages, 98.2 % - 100 % between G phages (including FCL-2) and 94.9 % - 100 % between C-phages. Notably, Cluster A-phages originating from different countries (Figure 6a), shared surprisingly high level of genetic identity. Within each phage cluster the hot spots for genetical variability were found in different parts of the genomes (compared with the consensus, Figure 6). In Cluster A-phages the hot spot was located in the area coding for putative replication proteins, while in C-phages it was in the area encoding putative tail proteins. In addition, the area around 25 kbp (from the genome start), which has been speculated to code for tail fiber proteins (e.g. in FCOV-F25 ORFs 35 and 36) (Laanto *et al.*, 2017), was also characterized by variability among phages.

**Figure 6.**
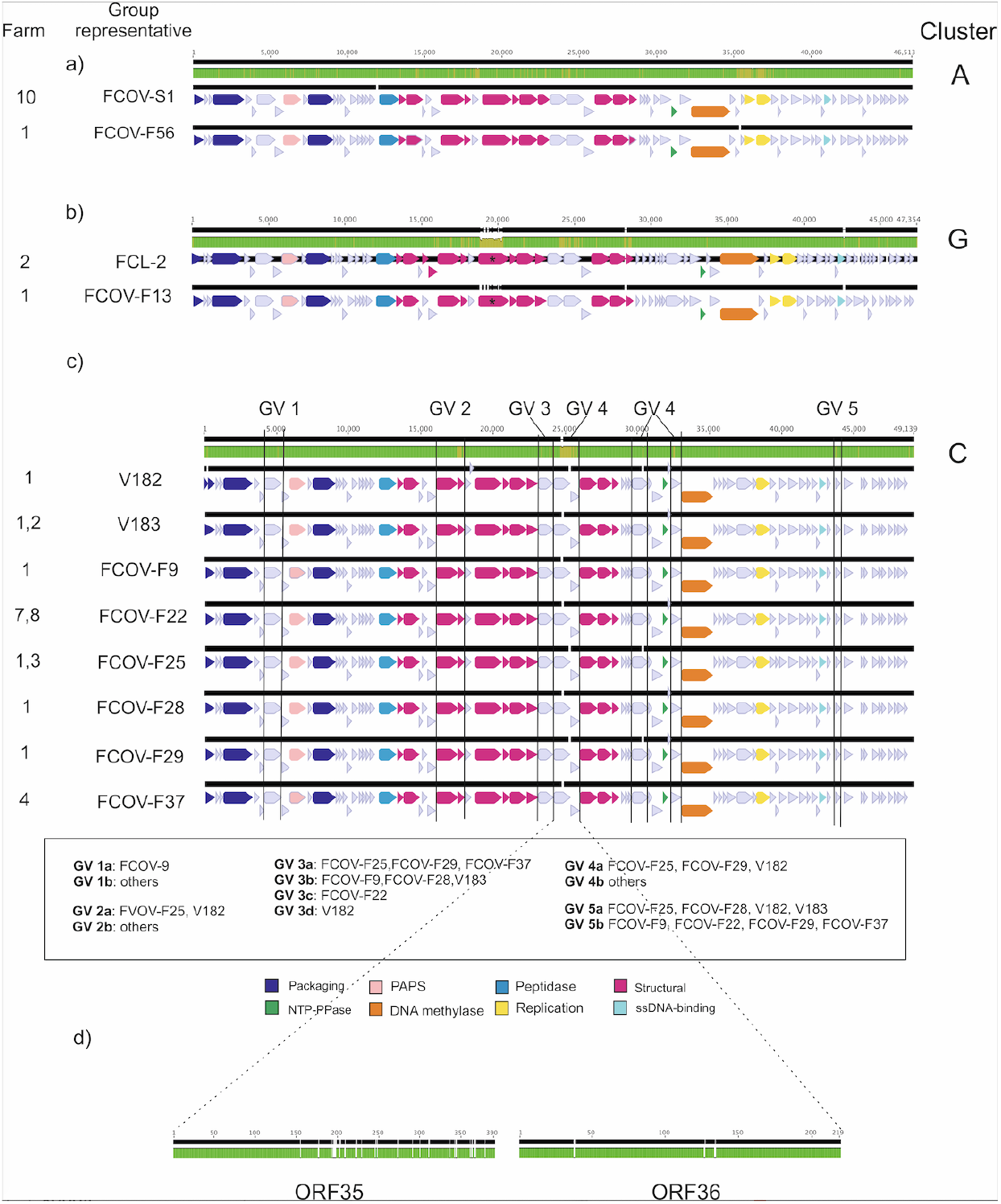
**Nucleotide alignments** of genetically different phages infecting *Flavobacterium columnare.* a) Cluster A (FVOV-S1 from Sweden and FCOV-F56 from Finland), b) Cluster G (FCL-2 isolated in 2008), and c) Cluster C phages (previously analysed V182 as a reference), and d) consensus of the amino acid alignments of ORF 35 and ORF 36 (speculated tail fiber proteins) between FCOV-F25 and FCOV-F28, which may be linked with differences in host range. In the genome consensus (on top of each alignment) green color indicates 100 % identical sequence and yellow >30 %. GV indicates genetically variable area (GV 1-5) where the differences localize. ORFs with unknown function are marked with grey, ORFs with putative function are marked with colors indicating the putative function as marked in the bottom. Letter after GV (a, b, c or d) indicate phages, which are similar in certain GV area but differ in other areas. PAPS = 3’-phosphoadenosine-5’-phosphosulfate. Asterix marks the tape tail measure protein in case of cluster G phages. Phage V182 isolated in 2014 was included in alignments as a reference to the last time point of the phage genome evolution dataset published earlier (Laanto et al., 2017).

Within Cluster G phages, only two nucleotide-level differences were detected in the phages FCOV-F45 and FCOV-F46 (same difference in both compared to others), one in non-coding area and one in a putative tail protein leading to amino acid change. Yet these phages differ in their host range, with FCOV-F46 able to infect also two C-genotype bacteria. The genome of the previously (in 2008) isolated phage FCL-2 was the only Cluster G phage to differ more from the other phages in that cluster. Interestingly, the putative tail tape measure protein of FCL-2 differs from the phages isolated in 2017 (Figure 6 b). Similarly to Cluster C phages, differences between FCL-2 and other G phages are seen in the ORF coding for speculated tail fiber proteins (hypothetical proteins before 25 kbp). Genetically identical phages are indicated in Table 1.

In general, the nucleotide level differences leading to amino acid level changes were detected in the putative structural proteins, in addition to several ORFs without putative annotated function (hypothetical proteins). Detailed list of differing ORFs is provided in Supplementary Table S5. In cluster A phages, changes were seen also in the putative 3’-phosphoadenosine-5’-phosphosulfate (PAPS), putative DNA methylase (ORF 47), and in the putative replication proteins (ORFs 50 and 51). In cluster G phages, the additional ORFs with amino acid changes were the putative peptidase (ORF 16) and ssDNA-binding protein (ORF 63).

Cluster C-phages were isolated most frequently, allowing higher molecular resolution of phage genomes in this group. Although the genomes were highly identical (Figure 5, Table 1), comparisons revealed seven genetically variable areas, forming five groups (GV 1-5) where nucleotide differences leading to amino acid change occurred (Figure 6c). The variation and similarity was not dependent on bacterial strain used in phage isolation (Table 1).

Most of the genetic variance occurred in the hypothetical proteins, but GV2 includes putative tail protein (ORF27). In addition, GV 4 includes the speculated tail fiber proteins in the 25 kbp area. Here, amino acid-level differences in ORFs 35 and 36 between FCOV-F25 and FCOV-F28 might explain differences in host range (Figure 6, Supplementary Table S4). It should also be noted that genetically identical phages were isolated from different fish farms, e.g. cluster C phages FCOV-F4 and F6 (from Farms 1 and 3, respectively) and FCOV-F20 and F-22 (from Farms 5 and 7), and Cluster G phages FCOV-F45 and −F54 (Farms 2 and 1) (Table 1).

## Discussion

In this study, we examined phenotypic and genetic characteristics of 126 *F. columnare* isolates and 63 phages from fish farms in Finland and Sweden. Bacterial isolates represented two previously characterized genomovars (I, I/II, based on 16S rDNA, (LaFrentz *et al.*, 2014), and four genetic groups (A, C, G, E, (Suomalainen *et al.*, 2006)). Bacteria belonging to genetic groups C and E had highest virulence in rainbow trout, but also other genetic groups caused high mortality. The isolated phages were all tailed *Myoviridae* dsDNA phages, and genetically similar compared to previously described *F. columnare* phages (Laanto *et al.*, 2017). The phages clustered into specific units of infection based on the bacterial genomic groups, with a few exceptions of phages able to cross infect to other bacterial groups (see below). Importantly, the isolated phages were able to impair the growth of the virulent bacteria, suggesting a potential to be used in phage therapy against columnaris disease.

Previous studies have shown that that genetically different *F. columnare* strains co-occur at fish farms (Ashrafi *et al.*, 2015; Sundberg *et al.*, 2016). Our results confirm the presence of previously isolated genetic groups in Finnish fish farms, suggesting that these bacterial populations have been circulating at farms during the last decades (Suomalainen *et al.*, 2006; Ashrafi *et al.*, 2015). Furthermore, the intensive aquaculture production in salmonid fish species in the Nordic countries may also select for certain host-associated *F. columnare* strains (LaFrentz *et al.*, 2018). Due to the convenient distance from the laboratory, Farm 1 was sampled the most frequently, and thus the majority of the isolated bacteria (51) and phages (30) were obtained from this farm, likely explaining the higher diversity obtained here. Nevertheless, the sampling showed presence of virulent *F. columnare* genetic groups (C and E) at almost all farms in Finland (Figure 1), and also in Sweden (exact farm locations for Swedish farms are not known). However, the Swedish *F. columnare* population was genetically more diverse with more genotypes isolated, some of which have not been found in Finland. Yet, the virulence of Swedish strains was lower than Finnish ones. To our knowledge, *F. columnare* isolates from Sweden have not been characterized previously.

We isolated phages infecting *F. columnare* from 6 farms in Finland and from one sample collected from Sweden. Interestingly, phages were also isolated when their bacterial hosts were not, suggesting that phages can be useful indicators of pathogen diversity during epidemics. Phage occurrence as a proxy for pathogen presence has been used also elsewhere in aquatic environments (Jofre *et al.*, 2016; McMinn *et al.*, 2017; Farkas *et al.*, 2019).

Phage infection patterns clustered according to *F. columnare* genetic groups, although there are a few exceptions. For example, although no phages were isolated using group E bacteria as isolation hosts, a few phages (FCOV-F25 – F27) had the ability to infect group E bacteria (FCO-F13 and −14) in addition to their isolation host (C). However, these phages did not genetically differ from other C-phages, which could only infect C type bacteria. Similarly, phage FCOV-F46 clustering to group G had the ability to infect a few group C bacteria, although being genetically identical to FCOV-F45 infecting only G bacteria. A previous study with Bacteroidetes phages showed similar results, as serial passage of phages in bacterial hosts can result in changes in host range, even without detectable genetic changes (Holmfeldt *et al.*, 2016). It is also possible that epigenetic modifications play a role in host range (Hattman, 2009) but that remains to be verified in these phages. However, some differences in host range were associated with clear genetic changes. FCOV-F28 was able to infect genetic groups C and G, and comparison to FCOV-F25 revealed several non-synonymous nucleotide changes in the previously speculated tail fiber genes, ORFs35 and 36 (Laanto *et al.*, 2017). In our previous study these same ORFs accumulated several mutations during a co-culture of *F. columnare* and phage FCV-1, leading to change in host range (Laanto *et al.*, 2020). Previous data with other species also suggests point mutations in tail fiber proteins increase phage infectivity (Uchiyama *et al.*, 2011; Boon *et al.*, 2020).

Although the increased accumulation of phage genomic and metagenomics data has revealed their enormous genetic diversity both on global and local scales (George P. C. Salmond and Fineran, 2015), specific phages infecting specific hosts have been isolated across large geographical areas (Kellogg *et al.*, 1995; Wolf *et al.*, 2003; Sonnenschein *et al.*, 2017), suggesting that some groups of closely related phages may have a worldwide distribution. An important finding in this study is that similar phages exist at different fish farms and countries, although some small genetic differences occur. In addition, the isolated phage genomes were highly similar to the previously described phages in our dataset, as also reported earlier with phages infecting the genetic group C bacteria (Laanto *et al.*, 2017). Comparable findings have been reported also in other phage-bacterium systems, for example with aquaculture pathogens *F. psychrophilum* and *Vibrio anquillarum* (Castillo and Middelboe, 2016; Kalatzis *et al.*, 2017), where genetically similar phages were widely distributed across large temporal and spatial scales.

Phages infecting flavobacterial species are known to regulate the genetic and phenotypic diversity of their bacterial hosts. As a co-evolutionary response, this should select for diversity also in the phage population. Yet, *F. columnare* phages with 100 % nucleotide identity were isolated from different fish farms and using enrichment hosts isolated in different years (Table 1). For example, Cluster C phages FCOV-F29 (isolated with B537 from 2013) and FCOV-F31 (isolated with FCO-F2 from 2017) are identical, as are also e.g. V186 (B245 from 2009) and FCOV-F48 (B537), and A-phages FCOV-S1 and −S2 (B534 from 2013 and B067 from 2007, respectively). A possible explanation for the low genetic diversity among *F. columnare* phages includes a potential for transfer of phages and bacteria between farms with the fish stocks or via water sources. This could explain the 100% similarity of phages from Farms 5 and 7, which are located close to each other and share the same source of water. Similarly, the shared water source and transfer of fish fry from Farm 1 to Farm 3 may contribute to the isolation of identical phages at these farms. On the other hand, A-phages originating from Finland and Sweden shared 99,7% identity despite the geographical distance and different isolation hosts. Therefore, the low genetic diversity and the high costs of phage resistance (Laanto *et al.*, 2012, 2014) in *F. columnare* may select for a low genetic diversity of the phages. Also, the use of antibiotics may play a central role in maintaining the low genetic diversity of the host bacteria in aquaculture. Application of antibiotic treatment to control disease epidemics rapidly impact the bacterial population size, leaving less possibilities for phage-bacterium interaction at short (within-season) time span.

When looking at the temporal differences of phage isolates our results are in accordance with previous findings: the phages isolated in most recently (2017) had broader host range than the previously isolated phages (FCV-1 (2008), V183-V189 (2015), Cluster C, Supplementary Table S4) from the same fish farm, which were, mostly able to infect isolates from earlier time points. This indicates a coevolutionary response to evolution of bacterial resistance towards previously isolated phages. At the same time, the bacterial isolates were susceptible to contemporary phages, which may have evolved to overcome the resistance mechanisms of the hosts. Similar results from environmental data has been derived also in other phage-bacterium systems (Koskella and Parr, 2015). However, the low diversity both in the phage and host *F. columnare* populations seem to restrict the genetic changes to small areas in the phage genomes. Our genetic data indicate changes at the end of the genome which might explain why host range between 2015 and 2017 phages differ in Cluster C phages. Phage V182 (isolated in 2014, Farm 1) used as a reference in genome alignment (Figure 6c) was distinct to 2015 phages from the same farm. V182 was the most recent phage isolate used in our previous study on phage-bacterium coevolution during 2007-2014 at Farm 1 (Laanto et al, 2017). The genetic comparisons thus suggest that bacterial resistance mechanisms cause for directional selection in the phage genome over long time spans. However, this phenomenon was not detectable in isolates obtained within one outbreak season at Farm 1, as phages isolated during three time points were identical (Table 1). This might have been caused by other factors at farms (e.g. the use of antibiotics) or in the natural waters. Similar results can be derived from comparison of Cluster G phages isolated from farm 2. FCL-2 (isolated in 2008) was genetically different to phages isolated in 2017, which can also be seen in differences in the host range.

In relation to the genetic similarity, another main finding of our data is that phages isolated from certain farms were able to infect bacterial hosts from other farms. This indicates that the aquatic farming environment (probably together with fish transfers and other reasons mentioned above) does not form isolation barriers between geographic locations, which would lead to locally adapting phage populations. At nucleotide level, some farm-specific differences between the phages were observed, but this did not impact their host range. In a broader perspective, none of the phages were able to infect *F. columnare* strains isolated from Central Europe or USA, and also in Swedish hosts the infectivity was limited to few strains. It is therefore likely that phage-bacterium coevolution has different trajectories in different geographic areas. This remains to be demonstrated until phage - *F. columnare* interactions have been characterized outside Nordic countries. In addition, further genomic analysis of the host bacteria could reveal the resistance mechanisms in bacteria, explaining the phage host range between farms and in time.

Because of specific fatal effect against bacteria, lytic phages have been considered and studied as antimicrobial agents against bacterial infections instead of antibiotics (Lin *et al.*, 2017; Ooi *et al.*, 2019). Here we characterized phage diversity against *F. columnare* and its four different genetic groups. In this study, we increased the collection of the isolated and characterized *F. columnare* strains and phages from different fish farms. Our findings suggest phages capable of infecting virulent *F. columnare* strains are present at fish farms and these phages could be used as potential antimicrobial agents in future applications.

## Acknowledgements

This work resulted from the BONUS FLAVOPHAGE project supported by BONUS (Art 185), funded jointly by the EU and Academy of Finland and Innovation Fund Denmark. We also acknowledge funding from Academy of Finland (grants #314939 and #321985) and Jane and Aatos Erkko Foundation.

The authors wish to thank Dr. Päivi Rintamäki (University of Oulu), Eva Jansson (National Veterinary Institute of Sweden), Mr Yrjö Lankinen and Natural research Institute Finland for donating water samples and bacterial isolates, and Ms. Hannah Kempf, Dr. Roghaieh Ashrafi and special laboratory technician Mr. Petri Papponen for technical assistance. We also thank Jean-Francois Bernardet (INRA, France), Mark McBride (UWM, USA) and Attila Karsi for providing bacterial strains for phage host range testing.

L.-R.S., G.M.F.A., E.L., and the University of Jyväskylä are responsible for a patent application covering the commercial use of purified mucin for production, quantification, and isolation of bacteriophages. It is titled “Improved methods and culture media for production, quantification and isolation of bacteriophages” and was filed with the Finnish Patent and Registration Office under patent no. FI20185086 (PCT/FI2019/050073) on 31 January 2018.

