## SUPPLEMENTARY INFORMATION for "Prevalence of genetically similar *Flavobacterium columnare* phages across aquaculture environments reveals a strong potential for pathogen control"

### Supplementary Figures

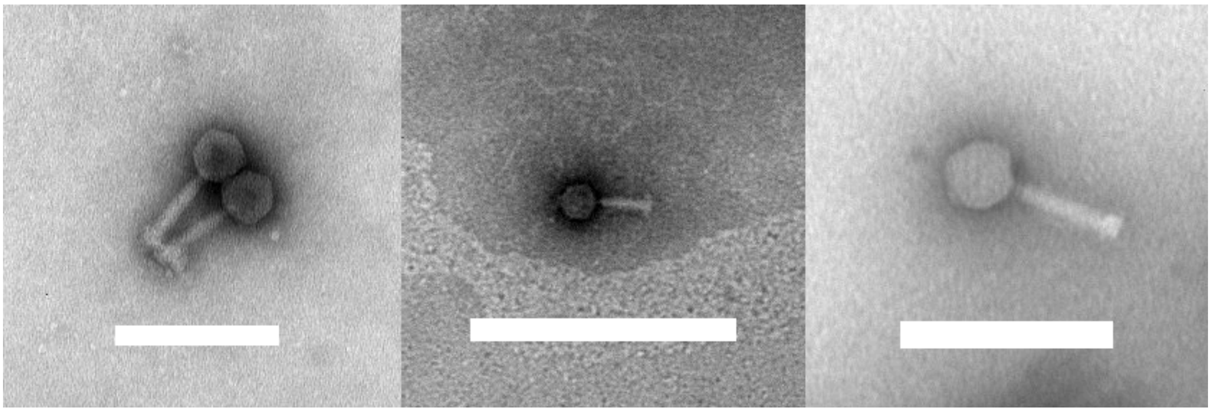

**Supplementary Figure S1.** Transmission electron microscopy (TEM) images of representative *F. columnare* phages infecting different genetic groups of the host (A, C and G). A). FCOV-S1, B). FCOV-F13 and C). FCOV-F27. Scale bar in A) and C) is 200 nm and B) 500 nm.

**Supplementary Table S1.** Bacterial strains isolated and/or characterized in this study. Swedish fish farms are confidential and not known by the authors, so they are jointly marked as Farm 10

| Isolate | Isolation year | Fish farm n:o | Fish species | Source | Genomovar | Genetic group | Virulence tested |
|---|---|---|---|---|---|---|---|
| F194 | 1997 | Sweden/10 | Brown trout | Kidney | I | E | |
| F195 | 1997 | Sweden/10 | Brown trout | Skin | I | A | |
| F310 | 2000 | Sweden/10 | Brown trout | Skin | I/II | ND3 | yes |
| F383 | 2002 | Sweden/10 | Salmon | Skin | I/II | ND3 | yes |
| F387 | 2002 | Sweden/10 | Brown trout | Skin | I | E | yes |
| F397 | 2002 | Sweden/10 | Brown trout | Kidney | I | ND1 | yes |
| F512 | 2013 | Sweden/10 | Brown trout | Kidney | I | ND2 | yes |
| F514 | 2013 | Sweden/10 | Brown trout | Kidney | I/II | ND3 | yes |
| F524 | 2014 | Sweden/10 | Rainbow trout | Kidney | I | A | yes |
| 3/3449 | 2017 | Sweden/10 | Rainbow trout | Skin ulcer | I | A | yes |
| 4/3450 | 2017 | Sweden/10 | Rainbow trout | Skin ulcer | I | A | |
| 5/3451 | 2017 | Sweden/10 | Rainbow trout | Skin ulcer | I | A | |
| 5/3460 | 2017 | Sweden/10 | Rainbow trout | Kidney and gills | I | A | |

| | | | | | | | |
|---|---|---|---|---|---|---|---|
| 6/3461 | 2017 | Sweden/10 | Rainbow trout | Kidney and gills | I | A | |
| FCO-S1 | 2017 | Sweden/10 | Rainbow trout | Fish tissue | I | A | yes |
| FCO-F1 | 2017 | 1 | Salmon | Tank water | I | C | |
| FCO-F2 | 2017 | 1 | Salmon | Tank water | I | C | yes |
| FCO-F3 | 2017 | 2 | Lake trout | Dorsal fin | I | G | yes |
| FCO-F4 | 2017 | 2 | Sea trout | Gill | I | G | |
| FCO-F5 | 2017 | 2 | Lake trout | Gill | I | G | yes |
| FCO-F6 | 2017 | 2 | Lake trout | Gill | I | G | |
| FCO-F7 | 2017 | 2 | Lake trout | Gill | I | G | |
| FCO-F8 | 2017 | 2 | Lake trout | Tank water | I | G | yes |
| FCO-F9 | 2017 | 2 | Sea trout | Tank water | I | G | yes |
| FCO-F10 | 2017 | 2 | Sea trout | Tank water | I | G | |
| FCO-F11 | 2017 | 3 | Rainbow trout | Gill | I | E | yes |
| FCO-F12 | 2017 | 3 | Rainbow trout | Gill | I | E | |
| FCO-F13 | 2017 | 3 | Rainbow trout | Tank water | I | E | yes |
| FCO-F14 | 2017 | 3 | Rainbow trout | Tank water | I | E | |
| FCO-F15 | 2017 | 3 | Rainbow trout | Tank water | I | E | |
| FCO-F16 | 2017 | 3 | | Outgoing water | I | E | yes |
| FCO-F19 | 2017 | 1 | Rainbow trout | Gill | I | C | |
| FCO-F20 | 2017 | 1 | Rainbow trout | Pectoral fin | I | C | |
| FCO-F21 | 2017 | 1 | Rainbow trout | Gill | I | C | |
| FCO-F22 | 2017 | 1 | Rainbow trout | Gill | I | C | yes |
| FCO-F23 | 2017 | 1 | Rainbow trout | Gill | I | C | |
| FCO-F24 | 2017 | 1 | Rainbow trout | Gill | I | C | |
| FCO-F25 | 2017 | 1 | Rainbow trout | Gill | I | C | |
| FCO-F26 | 2017 | 1 | Rainbow trout | Tank water | I | C | |
| FCO-F27 | 2017 | 1 | Rainbow trout | Tank water | I | C | |
| FCO-F28 | 2017 | 1 | Rainbow trout | Tank water | I | C | |
| FCO-F29 | 2017 | 1 | Rainbow trout | Tank water | I | C | |
| FCO-F30 | 2017 | 1 | Rainbow trout | Tank water | I | C | yes |
| FCO-F31 | 2017 | 1 | Rainbow trout | Tank water | I | C | |
| FCO-F32 | 2017 | 1 | Rainbow trout | Tank water | I | A | yes |
| FCO-F33 | 2017 | 1 | Rainbow trout | Tank water | I | C | yes |
| FCO-F35 | 2017 | 1 | | Outgoing water | I | C | |
| FCO-F36 | 2017 | 1 | | Outgoing water | I | C | |
| FCO-F37 | 2017 | 1 | | Outgoing water | I | C | |
| FCO-F40 | 2017 | 1 | | Outgoing water | I | C | yes |
| FCO-F41 | 2017 | 1 | | Outgoing water | I | C | yes |
| FCO-F42 | 2017 | 8 | Lake trout | Kidney | I | A | yes |
| FCO-F43 | 2017 | 8 | Lake trout | Gill | I | A | |
| FCO-F44 | 2017 | 8 | Lake trout | Kidney | I | A | |
| FCO-F45 | 2017 | 8 | Lake trout | Tank water | I | A | yes |
| FCO-F46 | 2017 | 8 | Lake trout | Tank water | I | A | |
| FCO-F47 | 2017 | 8 | | Outgoing water | I | A | yes |
| FCO-F49 | 2017 | 8 | | Outgoing water | I | A | |
| FCO-F50 | 2017 | 1 | Salmon | Gill | I | C | yes |
| FCO-F51 | 2017 | 1 | Salmon | Gill | I | C | |
| FCO-F52 | 2017 | 1 | Salmon | Gill | I | C | |

| | | | | | | | |
|---|---|---|---|---|---|---|---|
| FCO-F53 | 2017 | 1 | Salmon | Gill | I | C | |
| FCO-F54 | 2017 | 1 | Salmon | Gill | I | C | |
| FCO-F55 | 2017 | 1 | Salmon | Gill | I | C | |
| FCO-F56 | 2017 | 1 | Salmon | Gill | I | C | |
| FCO-F57 | 2017 | 1 | Salmon | Gill | I | C | |
| FCO-F58 | 2017 | 1 | Salmon | Tank water | I | C | yes |
| FCO-F59 | 2017 | 1 | Salmon | Tank water | I | C | |
| FCO-F60 | 2017 | 1 | Salmon | Tank water | I | C | |
| FCO-F61 | 2017 | 1 | Salmon | Tank water | I | C | |
| FCO-F62 | 2017 | 1 | Salmon | Tank water | I | C | |
| FCO-F63 | 2017 | 1 | Salmon | Tank water | I | C | |
| FCO-F64 | 2017 | 1 | Salmon | Tank water | I | C | |
| FCO-F65 | 2017 | 1 | Salmon | Tank water | I | C | |
| FCO-F66 | 2017 | 1 | Salmon | Tank water | I | C | |
| FCO-F67 | 2017 | 1 | Salmon | Tank water | I | C | |
| FCO-F68 | 2017 | 1 | Salmon | Tank water | I | C | |
| FCO-F69 | 2017 | 1 | Salmon | Tank water | I | C | |
| FCO-F70 | 2017 | 1 | Salmon | Tank water | I | C | |
| FCO-F71 | 2017 | 1 | Salmon | Tank water | I | C | |
| FCO-F72 | 2017 | 1 | Salmon | Tank water | I | C | |
| FCO-F73 | 2017 | 1 | Salmon | Tank water | I | C | |
| FCO-F74 | 2017 | 1 | Salmon | Tank water | I | C | |
| FCO-F75 | 2017 | 1 | Salmon | Tank water | I | C | |
| FCO-F76 | 2017 | 1 | Salmon | Tank water | I | C | |
| FCO-F77 | 2017 | 1 | Whitefish | Gill | I | C | |
| FCO-F78 | 2017 | 7 | Lake trout | Kidney | I | C | yes |
| FCO-F80 | 2017 | 7 | Lake trout | Pectoral fin | I | C | |
| FCO-F81 | 2017 | 6 | Salmon | Gill | I | E | yes |
| FCO-F82 | 2017 | 6 | Salmon | Kidney | I | E | |
| FCO-F83 | 2017 | 6 | Salmon | Kidney | I | E | |
| FCO-F84 | 2017 | 6 | Salmon | Kidney | I | E | |
| FCO-F85 | 2017 | 6 | Salmon | Kidney | I | E | |
| FCO-F86 | 2017 | 6 | Salmon | Gill | I | E | yes |
| FCO-F87 | 2017 | 1 | | Outgoing water | I | C | |
| FCO-F88 | 2017 | 6 | Salmon | Tank water | I | E | yes |
| FCO-F89 | 2017 | 6 | Salmon | Tank water | I | E | |
| FCO-F90 | 2017 | 6 | Salmon | Tank water | I | E | |
| FCO-F91 | 2017 | 6 | Salmon | Tank water | I | E | |
| FCO-F92 | 2017 | 6 | Salmon | Tank water | I | E | |
| FCO-F93 | 2017 | 6 | Salmon | Tank water | I | E | |
| FCO-F94 | 2017 | 6 | Salmon | Tank water | I | E | |
| FCO-F95 | 2017 | 6 | Salmon | Tank water | I | E | |
| FCO-F96 | 2017 | 6 | Salmon | Tank water | I | E | |
| FCO-F97 | 2017 | 6 | Salmon | Tank water | I | E | |
| FCO-F98 | 2017 | 5 | Sea trout | Skin/skin lesion | I | C | yes |
| FCO-F99 | 2017 | 5 | Sea trout | Skin/skin lesion | I | C | |
| FCO-F100 | 2017 | 5 | Sea trout | Skin/skin lesion | I | C | |
| FCO-F101 | 2017 | 5 | Sea trout | Skin/skin lesion | I | C | |

| | | | | | | | |
|---|---|---|---|---|---|---|---|
| FCO-F102 | 2017 | 5 | Sea trout | Skin/skin lesion | I | C | |
| FCO-F103 | 2017 | 5 | Sea trout | Skin/skin lesion | I | C | |
| FCO-F104 | 2017 | 5 | Sea trout | Skin/skin lesion | I | C | |
| FCO-F105 | 2017 | 5 | Sea trout | Skin/skin lesion | I | C | |
| FCO-F106 | 2017 | 5 | Sea trout | Skin/skin lesion | I | C | |
| FCO-F107 | 2017 | 5 | Sea trout | Skin/skin lesion | I | C | |
| FCO-F108 | 2017 | 5 | Sea trout | Skin/skin lesion | I | C | |
| FCO-F109 | 2017 | 5 | Sea trout | Skin/skin lesion | I | C | |
| FCO-F110 | 2017 | 5 | Sea trout | Skin/skin lesion | I | C | |
| FCO-F111 | 2017 | 5 | Sea trout | Skin/skin lesion | I | C | |
| FCO-F112 | 2017 | 5 | Sea trout | Skin/skin lesion | I | C | |
| FCO-F113 | 2017 | 5 | Sea trout | Skin/skin lesion | I | C | |
| FCO-F114 | 2017 | 5 | Sea trout | Skin/skin lesion | I | C | |
| FCO-F115 | 2017 | 5 | Sea trout | Skin/skin lesion | I | C | |
| FCO-F116 | 2017 | 5 | Sea trout | Skin/skin lesion | I | C | |
| FCO-F117 | 2017 | 5 | Sea trout | Skin/skin lesion | I | C | |
| FCO-F118 | 2017 | 7 | Salmon | Kidney | I | C | yes |

**Supplementary Table S2.** Previously isolated *F. columnare* bacterial strains used in host range studies. *F. columnare* strains from H to B533 have been previously characterized (Ashrafi *et al.*, 2015). Strains marked with asterisks were used in phage isolation. Strains collected from farms 5-9 were isolated and kindly donated by Dr. Päivi Rintamäki.

| **Bacterial strain** | **Genomovar** | **Genetic group** | **Isolation year** | **Isolation farm** | **Isolation source** |
|---|---|---|---|---|---|
| H | I | H | 2003 | 1 | rainbow trout (*Oncorhynchus mykiss*) |
| Tulo2 | I | A | 2010 | 1 | water |
| C1* | I | C | | unknown | |
| B067* | I | A | 2007 | 2 | Atlantic salmon (*Salmo salar*) |
| B185* | I | G | | 2 | water |
| B230 | I | C | 2009 | 1 | water |
| B245 | I | C | 2009 | 1 | water |
| B259 | I | C | 2010 | 1 | water |
| B261 | I | C | 2011 | 1 | water |
| B269 | I | A | 2009 | 1 | water |
| B270 | I | C | 2010 | 1 | water |
| B357 | I | C | 2010 | 1 | water/earthen pond |
| B366 | I | C | 2010 | 1 | farm oulet water |
| B367 | I | E | 2010 | 1 | water |
| B369 | I | E | 2010 | 1 | brown trout (*Salmo trutta*) |
| B376 | I | E | 2010 | 1 | water |
| B377 | I | E | 2010 | 1 | water/earthen pond |
| B379 | I | E | 2010 | 1 | farm oulet water |
| B393 | I | G | 2010 | nature/Hankasalmi | bream (*Abramis brama* ) |
| B396 | I | A | 2010 | 1 | inlet biofilm |
| B398 | I | A | 2010 | 1 | inlet water |

| | | | | | |
|---|---|---|---|---|---|
| B399 | I | G | 2010 | 1 | water |
| B402 | I | C | 2010 | 1 | European whitefish(*Coregonus lavaretus*) |
| B405 | I | C | 2010 | nature/Lake Jyvasjarvi | water |
| B407 | I | C | 2010 | nature/Hankasalmi | water |
| B408 | I | C | 2010 | nature/Hankasalmi | water |
| B409 | I | C | 2010 | 7 | sea trout (*Salmo trutta trutta*) |
| B416 | I | C | 2008 | 7 | brown trout (*Salmo trutta)* |
| B417 | I | C | 2008 | 7 | Atlantic salmon (*Salmo salar*) |
| B418 | I | C | 2009 | 7 | brown trout (*Salmo trutta*) |
| B419 | I | C | 2009 | 8 | Atlantic salmon (*Salmo salar*) |
| B420 | I | G | 2009 | 8 | Atlantic salmon (*Salmo salar*) |
| B421 | I | C | 2008 | 8 | Atlantic salmon (*Salmo salar*) |
| B422 | I | A | 2009 | 9 | sea trout (*Salmo trutta trutta*) |
| B423 | I | A | 2009 | 9 | lake trout (*Salmo trutta lacustris*) |
| B424 | I | C | 2007 | 7 | brown trout (*Salmo trutta*) |
| B426 | I | C | 2006 | 8 | Atlantic salmon (*Salmo salar)* |
| B427 | I | A | 2006 | 9 | brown trout (*Salmo trutta*) |
| B429 | I | H | 2003 | 2 | Pike perch (*Stizostedion lucioperca*) |
| B430 | I | E | 2003 | 2 | Pike perch (*Stizostedion lucioperca*) |
| B431 | I | A | 2003 | 2 | grayling *Thymallus thymallus* |
| B434 | I | G | 2005 | 6 | Atlantic salmon (*Salmo salar*) |
| B435 | I | G | 2005 | 6 | Atlantic salmon (*Salmo salar*) |
| B436 | I | G | 2006 | 6 | Atlantic salmon (*Salmo salar*) |
| B437 | I | C | 2006 | 8 | brown trout (*Salmo trutta*) |
| B438 | I | A | 2006 | 9 | rainbow trout (*Oncorhynchus mykiss*) |
| B439 | I | C | 2006 | 3 | rainbow trout (*Oncorhynchus mykiss*) |
| B440 | I | G | 2007 | 8 | Atlantic salmon (*Salmo salar*) |
| B441 | I | C | 2006 | 8 | rainbow trout (*Oncorhynchus mykiss*) |
| B442 | I | G | 2007 | 6 | Atlantic salmon (*Salmo salar*) |
| B444 | I | G | 2007 | 6 | Atlantic salmon (*Salmo salar*) |
| B445 | I | E | 2007 | nature/Hankasalmi | water |
| B446 | I | E | 2007 | nature/Hankasalmi | water |
| B447 | I | C | 2007 | nature/Hankasalmi | water |
| B448 | I | C | 2007 | nature/Lake Jyvasjarvi | water |
| B449 | I | C | 2007 | nature/Lake Jyvasjarvi | water |
| B450 | I | G | 2007 | 2 | Pike perch (*Stizostedion lucioperca*) |
| B451 | I | G | 2007 | 2 | Pike perch (*Stizostedion lucioperca*) |

| | | | | | |
|---|---|---|---|---|---|
| B452 | I | A | 2007 | 2 | Pike perch (*Stizostedion lucioperca*) |
| B453 | I | E | 2008 | 1 | rainbow trout (*Oncorhynchus mykiss*) |
| B454 | I | E | 2008 | 1 | rainbow trout (*Oncorhynchus mykiss*) |
| B458 | I | C | 2009 | 8 | brown trout (*Salmo trutta*) |
| B463 | I | C | 2011 | 3 | rainbow trout (*Oncorhynchus mykiss*) |
| B480* | I | E | 2012 | 1 | rainbow trout (*Oncorhynchus mykiss*) |
| B491 | I | E | 2012 | 1 | water |
| B496 | I | E | 2012 | 1 | water |
| B503 | I | E | 2012 | 1 | rainbow trout (*Oncorhynchus mykiss*) |
| B504 | I | E | 2012 | 1 | rainbow trout (*Oncorhynchus mykiss*) |
| B508 | I | E | 2012 | 1 | rainbow trout (*Oncorhynchus mykiss*) |
| B510 | I | E | 2012 | 1 | rainbow trout (*Oncorhynchus mykiss*) |
| B511 | I | E | 2012 | 1 | rainbow trout (*Oncorhynchus mykiss*) |
| B513 | I | E | 2012 | 1 | rainbow trout (*Oncorhynchus mykiss*) |
| B517 | I | E | 2011 | nature/Hankasalmi | water |
| B518 | I | C | 2006 | 8 | Atlantic salmon (*Salmo salar*) |
| B519 | I | C | 2006 | 7 | brown trout (*Salmo trutta*) |
| B520 | I | C | 2006 | 7 | Atlantic salmon (*Salmo salar*) |
| B521 | I | C | 2006 | 7 | Atlantic salmon (*Salmo salar*) |
| B523 | I | A | 2012 | 9 | brown trout (*Salmo trutta*) |
| B526 | I | C | 2012 | 6 | Atlantic salmon (*Salmo salar)* |
| B529 | I | C | 2012 | 6 | Atlantic salmon (*Salmo salar)* |
| B531 | I | C | 2012 | 7 | brown trout (*Salmo trutta*) |
| B532 | I | C | 2012 | 7 | brown trout (*Salmo trutta*) |
| B533 | I | H | 2012 | 8 | Atlantic salmon (*Salmo salar)* |
| B534* | I | A | 2013 | 1 | rainbow trout (*Oncorhynchus mykiss*) |
| B537* | I | C | 2013 | 1 | rainbow trout (*Oncorhynchus mykiss*) |

**Supplementary Table S3.** Previously isolated *F. columnare* strains from USA and France, and other bacterial species used in host range studies.

| **Strain/species** | **Genomovar** | **Isolation year** | **Provided by** | **Reference (if available)** |
|---|---|---|---|---|
| ATCC49513 | I | 1987 | Jean-Francois Bernardet, INRA, France | (Bernardet, 1989) |
| ATCC49512 | I | 1987 | ” | (Bernardet, 1989) |
| LDA 39 | I | | ” | |
| NCIMB2248T | I | | | (Bernardet and Grimont, 1989) |
| CSF258-10 (USA) | I | | Prof Mark McBride, University of Wisconsin Milwaukee, USA | (Evenhuis *et al.*, 2016, 2017) |

| | | | | |
|---|---|---|---|---|
| MSFC4 (USA) | I | | ” | (Evenhuis *et al.*, 2014; Bartelme *et al.*, 2018) |
| 90-106 (USA) | III | 1990 | Dr Attila Karsi | (Soto *et al.*, 2008) |
| L90-629 (USA) | III | 1990 | ” | (Soto *et al.*, 2008) |
| S03-579 (USA) | II/B | 2005 | ” | |
| S09-108 (USA) | II | 2009 | ” | |
| S09-157 (USA) | III | 2009 | ” | |
| S10-025 (USA) | III | 2010 | ” | |
| S10-239 (USA) | II | 2010 | ” | |
| C-069 (USA) | II | 2010 | ” | |
| C-074 (USA) | II | 2010 | ” | |
| CB10-151 (USA) | I | 2010 | ” | |
| Flavobacterium sp.B330 | | | | (Laanto, Mäntynen, *et al.*, 2017) |
| Flavobacterium sp.B183 | | | | (Laanto, Ravantti, *et al.*, 2017) |
| F. johnsoniae UW101 | | | | (McBride and Braun, 2004) |
| F.psychrophilum 950106-1/1 | | | | (Stenholm *et al.*, 2008) |
| F.psychrophilum MH1 | | | | (Castillo *et al.*, 2012) |

**Supplementary Table S4.** Host range of phages in *F. columnare* strains isolated in this study, in previously isolated *F. columnare* strains, and in other flavobacterial species. Black square indicates infection (clear lysis in three consequent phage dilutions), grey square indicates growth inhibition, and white square no effect (i.e. bacterial resistance). Each column represents a phage isolate and each row represents a host isolate.

(SEPARATE EXCEL FILE)

**Supplementary Table S5.** List of open reading frames (ORFs) that displayed differences between phage genomes in phages infecting genetic group C, A and G hosts.

**CUSTER C PHAGES**

| ORF nr in FCOV-F9 | Phages where the difference exists. Identical sequences within columns. | | | | Amino acid/reading frame change | Predicted function |
|---|---|---|---|---|---|---|
| **ORF01** | V182 | rest | | | yes | small subunit terminase |
| **ORF07** | FCOV-F9 | rest | | | | helicase |
| **ORF27** | V182 FCOV-F25 | rest | | | yes | tail protein (tail connector) |
| **ORF28** | V182 FCOV-F26 | rest | | | yes | structural protein |
| **ORF34** | V182 | FCOV-F25<br>FCOV-F29<br>FCOV-F37 | FCOV-F9<br>FCOV-F28<br>V183 | FCOV-F22 | yes | Hypothetical protein |
| **ORF35** | V182 FCOV-F25<br>FCOV-F29 | FCOV-F9<br>FCOV-F22<br>FCOV-F29<br>FCOV-F37<br>V183 | | | yes | Hypothetical protein (tail fiber?) |
| **ORF36** | V182 FCOV-F25<br>FCOV-F29 | FCOV-F9<br>FCOV-F22<br>FCOV-F29<br>FCOV-F37<br>V183 | | | yes | Hypothetical protein (tail fiber?) |
| **ORF43** | V182 FCOV-F25<br>FCOV-F29 | FCOV-F9<br>FCOV-F22<br>FCOV-F29<br>FCOV-F37<br>V183 | | | yes | Hypothetical protein |
| **ORF48** | V182 FCOV-F25<br>FCOV-F29 | FCOV-F9<br>FCOV-F22<br>FCOV-F29<br>FCOV-F37<br>V183 | | | yes | Hypothetical protein |
| **ORF68** | V182 V183 FCOV-F25<br>FCOV-F28 | FCOV-F9<br>FCOV-F22<br>FCOV-F29<br>FCOV-F37 | | | yes | ssDNA-binding protein |

**CUSTER A PHAGES**

| ORF nr in FCOV-S1 | Phages where the difference exists. Identical sequences within columns. | | Leads to amino acid change/change in reading frame | Predicted function |
|---|---|---|---|---|
| ORF04 | V186 FCOV-F56 | FCOV-S1<br>FCOV-S2 | | large subunit terminase |
| ORF05 | V186 FCOV-F56 | FCOV-S1<br>FCOV-S2 | | hypothetical protein |
| ORF06 | V186 FCOV-F56 | FCOV-S1<br>FCOV-S2 | | hypothetical protein |
| ORF09 | V186 FCOV-F57 | FCOV-S1<br>FCOV-S2 | yes | phosphoadenosine phosphosulfate reductase family protein |
| ORF10 | V186 FCOV-F57 | FCOV-S1<br>FCOV-S2 | yes | hypothetical protein |
| ORF11 | V186 FCOV-F57 | FCOV-S1<br>FCOV-S2 | yes | portal protein |
| ORF12 | V186 FCOV-F57 | FCOV-S1<br>FCOV-S2 | | hypothetical protein |
| ORF22 | V186 FCOV-F57 | FCOV-S1<br>FCOV-S2 | yes | head decoration protein |
| ORF23 | V186 FCOV-F57 | FCOV-S1<br>FCOV-S2 | yes | major capsid protein |
| ORF24 | V186 FCOV-F57 | FCOV-S1<br>FCOV-S2 | yes | hypothetical protein |
| ORF27 | V186 FCOV-F57 | FCOV-S1<br>FCOV-S2 | yes | tail protein |
| ORF28 | V186 FCOV-F57 | FCOV-S1<br>FCOV-S2 | | tail protein |
| ORF29 | V186 FCOV-F57 | FCOV-S1<br>FCOV-S2 | yes | hypothetical protein |
| ORF30 | V186 FCOV-F57 | FCOV-S1<br>FCOV-S2 | yes | phage tape tail measure protein |
| ORF31 | V186 FCOV-F57 | FCOV-S1<br>FCOV-S2 | yes | hypothetical protein |
| ORF32 | V186 FCOV-F57 | FCOV-S1<br>FCOV-S2 | yes | baseplate protein |
| ORF33 | V186 FCOV-F57 | FCOV-S1<br>FCOV-S2 | yes | hypothetical protein |
| ORF34 | V186 FCOV-F57 | FCOV-S1<br>FCOV-S2 | yes | hypothetical protein |
| ORF35 | V186 FCOV-F57 | FCOV-S1<br>FCOV-S2 | yes | Hypothetical protein (tail fiber?) |
| ORF40 | V186 FCOV-F57 | FCOV-S1<br>FCOV-S2 | yes | hypothetical protein |
| ORF48 | V186 FCOV-F57 | FCOV-S1<br>FCOV-S2 | yes | hypothetical protein |
| ORF49 | V186 FCOV-F57 | FCOV-S1<br>FCOV-S2 | yes | DNA-methylase |
| ORF51 | V186 FCOV-F57 | FCOV-S1<br>FCOV-S2 | yes | hypothetical protein |
| ORF52 | V186 FCOV-F57 | FCOV-S1<br>FCOV-S2 | yes | hypothetical protein |
| ORF53 | V186 FCOV-F57 | FCOV-S1<br>FCOV-S2 | yes | replication |
| ORF54 | V186 FCOV-F57 | FCOV-S1<br>FCOV-S2 | yes | replicative helicase |
| ORF55 | V186 FCOV-F57 | FCOV-S1<br>FCOV-S2 | | hypothetical protein |
| ORF57 | V186 FCOV-F57 | FCOV-S1<br>FCOV-S2 | yes | hypothetical protein |
| ORF74 | V186 FCOV-F57 | FCOV-S1<br>FCOV-S2 | yes | |

**CUSTER G PHAGES**

| ORF nr in FCOV-F13 | Phages where the difference exists. Identical sequences within columns. | | | Leads to amino acid change/change in reading frame | Predicted function | Comments |
|---|---|---|---|---|---|---|
| ORF01 | FCL-2 | rest | | yes | small subunit terminase | |
| ORF13 | FCL-2 | rest | | yes | hypothetical protein | |
| ORF16 | FCL-2 | rest | | yes | peptidase | |
| ORF26 | FCL-2 | rest | | yes | tail protein | |
| ORF27 | FCL-2 | FCOV-F13<br>FCOV-F16<br>FCOV-F54 | FCOV-F45<br>FCOV-F46 | yes | tail seath protein | One amino acid change in FCOV-F45 and FCOV-F46 that is identical with FCL-2 |
| ORF28 | FCL-2 | rest | | yes | structural protein | |
| ORF29 | FCL-2 | rest | | yes | hypothetical protein | |
| ORF30 | FCL-2 | rest | | yes | tail tape measure protein | |
| ORF32 | FCL-2 | rest | | yes | baseplate protein | |
| ORF34 | FCL-2 | rest | | yes | hypothetical protein | |
| ORF35 | FCL-2 | rest | | yes | Hypothetical protein (tail fiber?) | |
| ORF36 | FCL-2 | rest | | yes | Hypothetical protein (tail fiber?) | |
| ORF37 | FCL-2 | rest | | | hypothetical protein | |
| ORF38 | FCL-2 | rest | | yes | tail protein | |
| ORF39 | FCL-2 | rest | | yes | structural protein | |
| ORF63 | FCL-2 | rest | | yes | ssDNA-binding | |
| ORF65 | FCL-2 | rest | | yes | hypothetical protein | |
| ORF70 | FCL-2 | rest | | yes | hypothetical protein | no ORF in FCL-2 |
